# Helicase: Vectorized parsing and bitpacking of genomic sequences

**DOI:** 10.64898/2026.03.19.712912

**Authors:** Igor Martayan, Loup Lobet, Camille Marchet, Charles Paperman

## Abstract

Modern sequencing pipelines routinely produce billions of reads, yet the dominant storage formats (FASTQ and FASTA) are text-based and sequential, making high-throughput parsing a persistent bottleneck in bioinformatics. Their regular, line-oriented structure makes them well-suited to SIMD vectorization, but existing libraries do not fully exploit it.

We present vectorized algorithms for high-throughput FASTA/Q parsing, with on-the-fly handling of non-ACTG characters and built-in bitpacking of DNA sequences into multiple compact representations. The parsing logic is expressed as a finite state machine, compiled into efficient SIMD programs targeting both x86 and ARM CPUs. These algorithms are implemented in *Helicase*, a Rust library exposing a tunable interface that retrieves only caller-requested fields, minimizing unnecessary work. Exhaustive benchmarks across a wide range of CPUs show that Helicase meets or exceeds the throughput of all evaluated state-of-the-art libraries, making it the fastest general-purpose FASTA/Q parser to our knowledge.

**Availability:** https://github.com/imartayan/helicase.

## 1 Introduction

Sequence bioinformatics has always been closely tied to stringology and compression theory, but connections to algebraic automata have remained comparatively rare. Recent progress has opened new paths toward high-performance parsers that are both formally grounded and practically competitive. This article explores one such connection, applying these techniques to the FASTA and FASTQ formats.

FASTA and FASTQ are among the oldest and most enduring data formats in bioinformatics. FASTA was introduced in 1985 alongside the sequence alignment tool of the same name [9,15], while FASTQ emerged in the early 2000s [2] as sequencing technologies began producing per-base quality scores. Over the years, both formats achieved near-universal adoption across the bioinformatics ecosystem. Nowadays, sequencing pipelines regularly produce billions of reads, and the overwhelming majority of this data is still stored and exchanged in FASTA and FASTQ. Public sequencing archives such as the European Nucleotide Archive (ENA) now contain hundreds of petabytes of sequencing data [3], meaning that infrastructure built around these text-based formats must now operate at scales their designers could not have anticipated. This results in a tension between the simplicity that made these formats successful and the performance demands of modern high-throughput bioinformatics.

### Description of the formats

FASTA and FASTQ are text-based formats containing a sequence of *records*. FASTA records contain two fields: (1) a single-line header starting with `‘>‘` followed by an optional description and (2) a multi-line ASCII-encoded sequence. The ASCII-encoded sequence contains `A/C/T/G` for DNA and other IUPAC codes to indicate ambiguous bases, and can be split into an arbitrary number of lines until the next record. A FASTA record can be matched with the following regular expression:

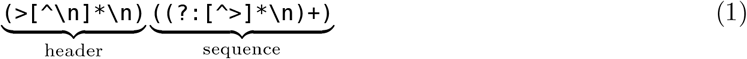

FASTQ records contain four fields: (1) a single-line header starting with ‘@’ followed by an identifier and an optional description, (2) a single-line ASCII-encoded sequence, (3) a single-line separator starting with `‘+’` optionally followed by an identifier and (4) a quality line of the same length of the sequence encoding the quality score of each base at the corresponding position. Compared to FASTA, the ASCII-encoded sequence also contains `A/C/T/G` and other IUPAC codes, but cannot be split into multiple lines. The quality line, however, may contain *any character* in the ASCII range [33, 126] including ‘@’, `‘+’` or any alphabetic character. Also note that, while the quality represents almost 50% of the content of a FASTQ file, it is often discarded during the analysis.

### Efficient parsing

At its core, parsing FASTA and FASTQ files reduces to a lexical analysis problem over large byte streams. The parser must identify structural delimiters, such as line feeds (`‘\n’`), header markers (`‘>‘` and ‘@’), and separators (`‘+’`) and segment the input into records accordingly. Since modern datasets often reach billions of reads, the dominant cost is no longer algorithmic complexity but the raw throughput at which these critical positions can be detected.

Reaching a high throughput requires minimizing branching and favoring regular, predictable computation patterns that can be efficiently executed on wide data paths. This constraint naturally shifts the focus from traditional parsing strategies toward approaches that process the input stream in bulk while limiting branches.

### Vectorization

Since the 1980s [17], CPUs have exposed SIMD (Single Instruction, Multiple Data) instructions that operate on wide registers, processing multiple bytes in parallel within a single instruction. Compilers can exploit these instructions automatically through auto-vectorization, and this proved to be effective for scientific and high-performance computing. However, auto-vectorization often fails when branches depend on input data, which is precisely the situation arising when parsing. Achieving high throughput therefore requires *manual* vectorization by encoding the parsing logic with SIMD intrinsics. A canonical example of a hand-optimized SIMD routine is `memchr`, which scans a memory region for the first occurrence of a given byte. While naive scalar implementations only reach a few GB/s, carefully vectorized ones approach the full RAM bandwidth at close to 40 GB/s on modern hardware. Parsing FASTA/Q, however, requires searching for *multiple* markers simultaneously (line feeds, header markers and separators) so relying on independent `memchr` calls leaves performance on the table. Fusing these searches into a single vectorized pass is the key algorithmic challenge we address in this contribution.

### Existing work

Most state-of-the-art parsers share a pretty similar design. The input file is first split into sequence of lines using `memchr` to locate line feeds (`‘\n’`). For FASTQ files, counting the lines is sufficient for the parsing logic since each field spans over a single line. For FASTA files, however, we also need to check the first character of the line to determine if it starts a header (`‘>‘`). This usually incurs a second check after each `memchr` match to detect a new record. In particular, both `kseq` [7] and `needletail` [12], two of the most popular parsers in the community, rely on this approach. According to Heng Li’s `biofast` benchmark [8], `needletail` is the fastest single-threaded parser currently available, so this will be the baseline we compare against.

Recent methods focused more specifically on parallelization strategies for parsing [22] and decompressing [14] FASTA/Q inputs. Among these works, RabbitFX [22] stands out as the state-of-the-art for parallel parsing of plain and gzipped FASTA/Q. However, since our work focuses on accelerating the parsing algorithm for a given thread, we will not evaluate RabbitFX in this article.

### Contributions

In this article, we present new vectorized algorithms for high-throughput FASTA/Q parsing, which support on-the-fly detection and handling non-ACTG characters, and provide bitpacked representations of the sequence. Formally, given an input FASTA/Q file, we produce a structured iterator over the (possibly bitpacked) content of each record. This approach is implemented in *Helicase*, a Rust library providing a configurable interface to expose caller-requested fields for each record, avoiding unnecessary computation. We evaluate our approach on both x86 and ARM architectures. Across all tested scenarios, Helicase matches or exceeds the throughput of existing single-threaded parsers while maintaining feature parity. In addition, it provides extended functionality, such as bitpacked sequence output, without compromising performance. For data already loaded in RAM, Helicase is able to parse uncompressed short reads at 27 GB/s and long reads at 49 GB/s on an Apple M3 Pro using a single thread, reaching the core memory bandwidth. Our library is available at https://github.com/imartayan/helicase under a permissive open-source license.

## 2 Methods

### 2.1 DNA representations

Depending on the application use case, different DNA representations might be desirable. Our approach produces three main representations: the traditional ASCII-based one, and two bitpacked versions (*packed* and *columnar*) in which each base is encoded with 2 bits. A convenient way to encode bases is to extract the second and third lowest bits from their ASCII representation, as described below:

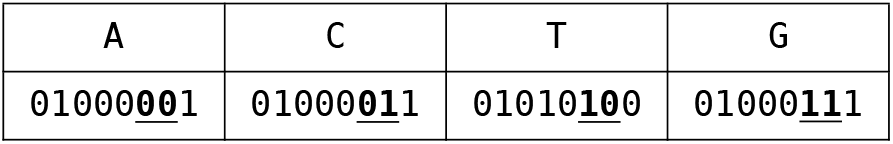

However, sequences may still contain non-ACTG characters for IUPAC codes that we cannot bitpack losslessly, so we have to handle them separately. We propose two different ways to solve this. One solution is to treat non-ACTG characters as splits and only return contiguous chunks of ACTGs. Another solution a lossy encoding that returns an additional bitmask marking ambiguous positions with ones (thus requiring 3 bits per base in total).

#### Packed format

Once each base is mapped to two bits, an intuitive way to represent the sequence is to pack each pair of bits consecutively, thus storing 4 bases in one byte. This bitpacked representation is probably the most commonly used, and allows simple iteration on the bases by shifting and masking.

#### Columnar format

Another way to represent the sequence is to separate the stream of high bits and low bits. With this approach, 8 bases are stored in two bytes: one for the high bits and the other for the low bits. This representation maps each base of the sequence to exactly one bit in each part at the corresponding position, which is convenient for bitmask-based operations. For instance, locating `T`s in the sequence is as simple as intersecting the high bits and the negation of the low bits:

| seq | G | A | T | T | A | C | A | T |
| --- | --- | --- | --- | --- | --- | --- | --- | --- |
| hi | 1 | 0 | 1 | 1 | 0 | 0 | 0 | 1 |
| lo | 1 | 0 | 0 | 0 | 0 | 1 | 0 | 0 |
| $hi \wedge \neg lo$ | 0 | 0 | <u>1</u> | <u>1</u> | 0 | 0 | 0 | <u>1</u> |

### 2.2 Streaming the input

Our approach follows three phases of *streaming, lexing* and *parsing*: the input is first streamed to a lexer that provides a tokenization of the stream, which is then fed to a parser to build the desired output.

The very first step of our algorithm is to stream the input that we want to parse. We support two main kinds of input: the ones that support random access (e.g. data in RAM or memory-mapped file) and the ones that have to be consumed sequentially (e.g. file reader or stdin). In particular, reader-based inputs allow transparent decompression of the data by detecting compressor-specific signatures (“magic bytes”) at the start of the input. To handle all these types of input generically, we provide a unifying abstraction that streams the data by blocks of fixed size (in our case 64 bytes). While streaming these blocks is straightforward for data in memory and does not require any copy, we need a buffering strategy for reader-based input to amortize the cost of read syscalls.

The buffering strategy differs depending on the type of output produced. For bitpacked representations, the input buffer is only needed for tokenization so we simply use a fixed-size buffer (128 KB by default) and refill it when it is entirely consumed. For the ASCII-based representation, however, we want to reuse the sequence loaded in the buffer whenever possible and avoid copies. In that case, we use an adaptive strategy similar to Needletail [12] or RabbitFX [22]. When a record is too large to fit in the current buffer, the size of the buffer is doubled. In practice, this occurs only for very long sequences such as chromosomes. When a record overlaps the boundary between two buffer fills, the partial record at the end of the current buffer is copied to the beginning of the next one before refilling.

### 2.3 Classifying the input with bitmasks

Unlike traditional lexing/parsing situations, we need to carefully design the structure of the lexing phase to be fully compatible with vectorized processing. Instead of consuming the input and producing lexemes to be consume later by the parser, the lexing phase will annotate the stream by producing bitmasks that encompass some information about the input. For instance, we will produce a bitmask that distinguishes parts of the input that are within the header of a FASTA record or within the sequence.

Producing bitmasks efficiently to classify the input is a classic way to produce highly efficient code. For instance, in simdjson [6], the lexing phase produces bitmasks for the position of structural characters in a JSON document. In rsonpath [4], the small programs generating relevant bitmasks are denoted *classifiers*, and we will follow this terminology from now on.

Concretely, a bitmask gives a boolean information for each byte of the input. Those bytes are provided in streaming by blocks of 64 at a time. The goal of a highly performant lexer is to produce the desired bitmasks as efficiently as possible, and with the minimal amount of branches. To illustrate this methodology, let us focus on the `header` bitmask, critical for the FASTA parsing in Helicase. This bitmask is true for a byte if and only if it belongs to a header of the document. Formally, it is delimited by the byte `‘>‘` and a line feed without any line feed in between, as described in the regular expression (1).

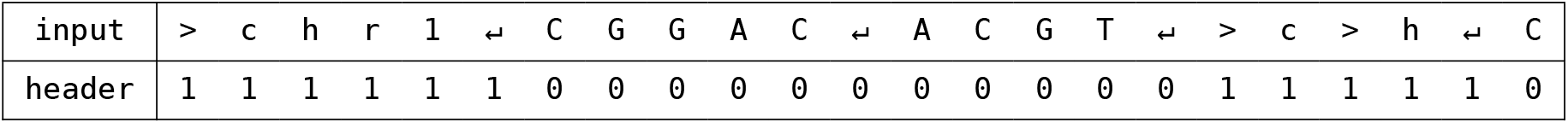

To construct such a bitmask, it is essential to avoid scanning the input sequentially one byte at a time. The first step is to identify the relevant byte values in the input stream. Following the methodology used in `memchr`, we can efficiently produce bitmasks (with architecture-specific implementations discussed later) that mark the occurrences of the characters `‘>‘` and `‘`↵’.

The goal is then to derive a bitmask that identifies precisely the bytes located between a `‘>‘` and a ‘↵’. Remarkably, this can be achieved without iterating over the bytes individually. Instead, the construction relies on carry propagation during the addition of two bit-vectors. This approach is efficient because modern processors perform additions on 64-bit integers with highly optimized carry propagation. In particular, the processor’s carry flag enables fast propagation of carries across successive additions, allowing the required bitmask to be computed with only a few arithmetic and bitwise operations.

#### From automata to vectorized programs

It is possible, using the work of [13,20,19], to analyze a given automaton and determine whether its execution can be expressed solely using bitwise bit-vector operations combined with addition. An automaton that admits an efficient program has several characterizations and belongs to the class of so-called *counter-free automata* [10]. Although these characterizations are decidable and conceptually well understood, there is currently no efficient implementation that automatically derives the corresponding vectorized program from the automaton. For small enough instances such as our lexer, the proof methodology of [13] can be applied manually to synthesize such a program. We can produce a compact representation of the lexing phase through so called *vectorial circuits*.

Those circuits can be thought of as formulas on bitmasks that can be evaluated on a given word. The results of the bitmask are the lexing annotations we need. Given a word *u* that we want to parse, we denote by 1_>_ and 1_↵_ the bitmask of position where respectively the symbols `‘>‘` and ‘↵’ occur. The formula to compute the `header` bitmask is then expressed as follows, using + to denote an addition propagating the carry from left to right, ¬ to denote bitwise `not` and ⊕ to denote bitwise `xor`:

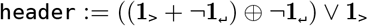

The usage of carry propagation to encode part of the branching originates from Myers [11]. Applying this formula to the previous example, we get the following:

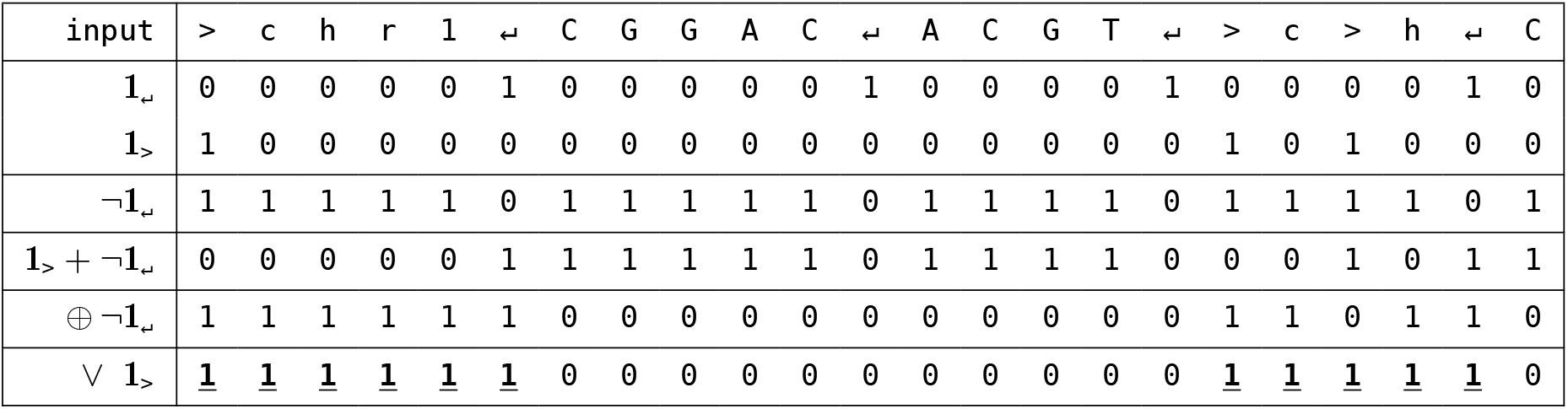

### 2.4 Implementation of the lexing phase

While this formula provides us with the appropriate semantics, implementing it requires a bit more work since it is interpreted over a stream of 64-bytes blocks. We now describe its concrete implementation in the lexing phase as a *classifier*, i.e. an iterator that builds the annotation on-the-fly.

The first step of the lexing phase is to load the input block into a SIMD register, and to compute bitmasks for specific characters: `‘\n’` for FASTQ, and both `‘\n’` and ‘>‘ for FASTA. Additionally, we may want to detect non-ACTG bases to split the sequence or mark their positions. Both of these steps are described in Algorithm 1 and rely essentially on vectorized byte-wise comparisons followed by a `movemask`. The `movemask` instruction, natively supported on x86 since SSE, extracts the most significant bit of each byte in a SIMD register into a scalar bitmask. However, this instruction is not available on ARM CPUs, so we have to implement it manually using multiple intrinsics, making it more expensive than on x86. A relatively efficient method to implement it on ARM is to make use of NEON’s interleaved layout and byte-wise packing^3^. One benefit of having a manual implementation is that we can easily adapt it to extract multiple bits from each byte instead of a single one, which will turn out to be useful for the packed representation.

#### Algorithm 1

Pseudocode to detect newlines and non-ACTG characters.

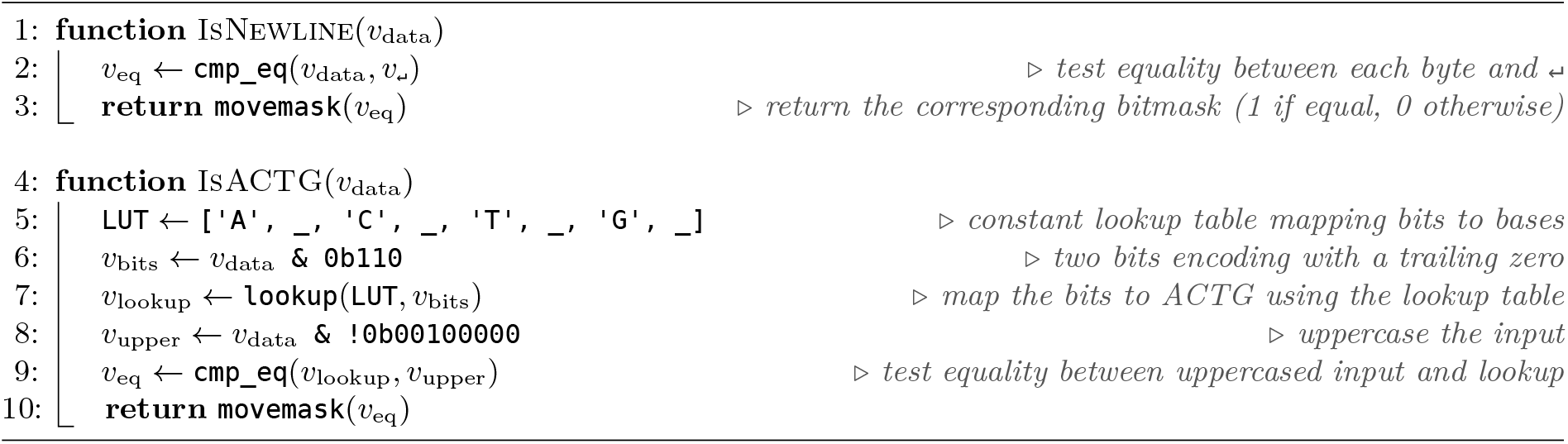

Depending on the type of desired output, we also use the lexing phase to produce a stream of columnar or packed sequences, both described in Algorithm 2. For the columnar representation, we simply shift each bytes by 5 (resp. 6) positions to move the high (resp. low) bit to the beginning of the byte, which we can then extract using `movemask` as before. The packed representation, however, requires different strategies depending on the architecture. On x86, since `movemask` can only extract one bit at a time, we essentially reuse the high and low bits from the columnar format and interleave them. A simple way to interleave the bits is to use the PDEP instruction available with BMI2. However, this instruction is not supported by old CPUs, and has been notoriously slow on AMD CPUs until the Zen3 generation because it was microcoded. As a fallback for this kind of hardware, we also provide an alternative implementation that does the interleaving step on the *bytes* before the `movemask` rather than the bits. On the other hand, computing packed representations on ARM is much simpler since we can change the `movemask` implementation to extract two bits at a time, thus entirely avoiding the interleaving step. Due to these architectural differences, producing a packed representation is slightly more expensive than a columnar one on x86.

#### Algorithm 2

Pseudocode to compute the columnar and packed representations, on x86 and ARM.

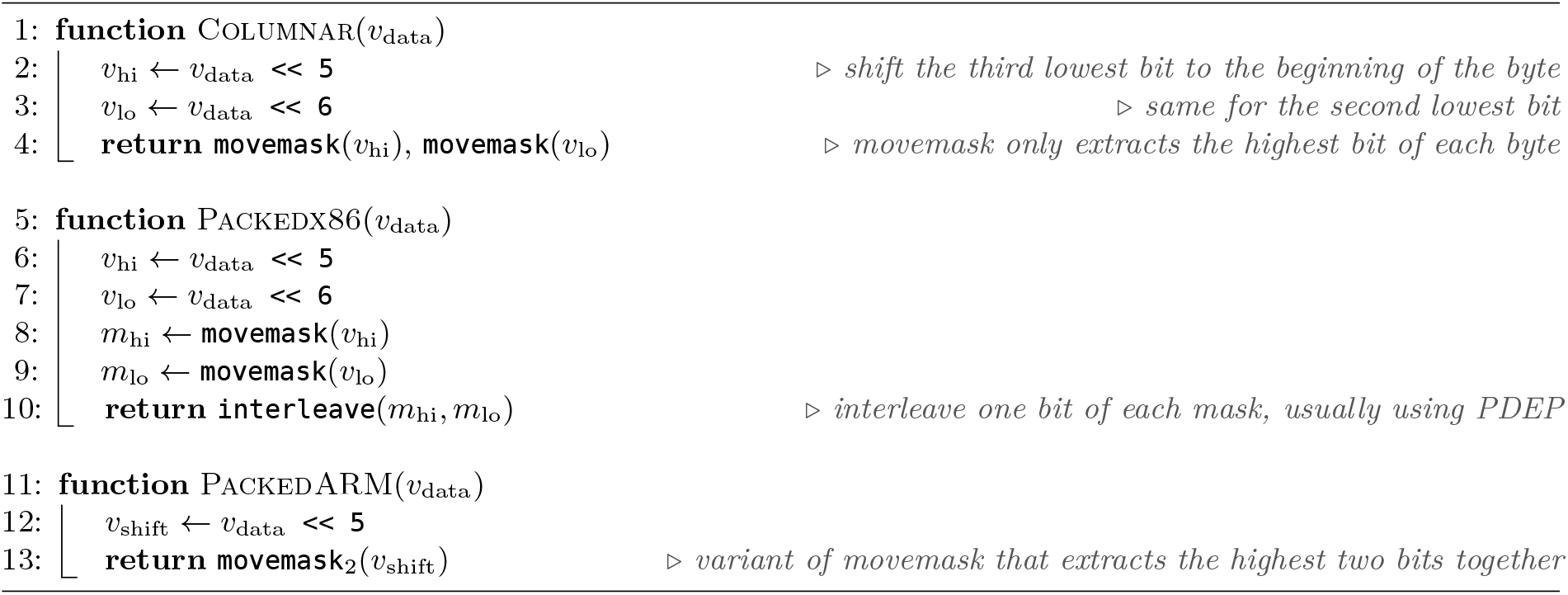

### 2.5 Parsing: extracting relevant information with a control finite state machine

The parsing stage is implemented as a deterministic control layer operating over the output of the lexing classifiers. Rather than inspecting input bytes individually, the parser consumes bitmask summaries associated with fixed-size blocks, where each mask encodes the positions of structurally relevant symbols such as header markers and DNA characters. This design allows the parser to advance directly between meaningful positions using bitwise selection, avoiding per-character control flow.

The control logic for FASTA parsing is structured around a small number of states corresponding to the successive processing steps. Execution begins by locating the start of a header, after which the header line is consumed. The parser then enters an intermediate phase where it determines whether the subsequent region corresponds to another header or to DNA content. When a DNA region is detected, the parser initializes the necessary context and enters a tight processing loop over contiguous DNA symbols (restricted to `A/C/T/G` for some bitpacking strategies). This loop continues as long as the lexical masks indicate valid DNA positions. Upon encountering a boundary condition, such as a non-DNA symbol, a header marker, or the end of the input, the current DNA segment is finalized and control returns to the intermediate phase. On FASTQ, the structure is similar but since each field spans over a single line, the corresponding states directly correspond to a given line count modulo 4, making it simpler overall.

All transitions are driven by predicates evaluated over the lexical masks. In practice, the parser repeatedly queries these masks to locate the next relevant position, using bitwise operations to identify the first occurrence of a header, DNA symbol or boundary. If no such position exists within the current block, the parser advances to the next block and repeats the process. As a result, large regions of unnecessary fields are skipped in constant time per block, and control decisions are made only at structurally significant locations.

The parser maintains a mutable state associated with the current record and DNA segment. Depending on the configuration, this state may include ranges for header and quality data, accumulators for DNA sequences, or alternative encodings such as packed and columnar representations. These data structures are updated incrementally during DNA processing and may be reset or extended when transitioning between segments. Boundary conditions determine when a DNA segment is considered complete, and user-defined policies control whether segments are split, merged or filtered.

The parsing stage can also produce events corresponding to record boundaries or DNA segments. These events are emitted at well-defined points in the control flow, typically when a new header is encountered or when a DNA segment is finalized. Event generation is entirely configuration-dependent and does not affect the structure of the control logic.

Overall, the parser follows a two-level organization in which a data-parallel lexing stage exposes structural information as bitmasks, and a compact control layer reacts to this information through a small set of states. This separation enables high-throughput streaming by eliminating fine-grained branching while preserving a simple and predictable execution model.

### 2.6 Compile-time configuration and specialization

The parser’s behavior is entirely dictated by a set of configuration parameters processed at compile time. These parameters control header processing, DNA accumulation strategy (string, packed, and/or columnar), boundary handling, and event emission. Conceptually, this design can be understood as generating a distinct parser implementation for each configuration, rather than a single generic parser containing runtime conditionals.

This approach corresponds to compile-time specialization (often referred to as “monomorphization”), where a generic implementation is transformed into multiple concrete variants. In contrast to designs relying on runtime branching (e.g., `if (compute_dna_columnar) …`), each generated variant contains only the code relevant to the selected configuration, with all other paths removed during compilation.

This specialization extends across both the parsing and lexing phases. Critical functions are aggressively inlined, and inlining propagates into the lexer itself. As a consequence, lexical computations that are not required under a given configuration are eliminated at compile time. For example, when DNA string accumulation is disabled, the corresponding logic for extracting and storing DNA bases is not generated, and the lexer omits the associated processing entirely. All the conditions governing buffer management, boundary semantics, and event emission are thus resolved statically. Each configuration hence produces a specialized instance of the control automaton in which transitions and associated actions are fixed in advance, with no residual dynamic branching. At the moment of writing, we provide 13 flags that can each trigger different behaviors. Altogether, this amounts to 8192 different possible specializations. More specialization could occur in the future to open the way for more efficient code.

## 3 Results

To evaluate the performance of our approach, we compared it against two state-of-the-art Rust libraries: Needletail 0.6.3 [12] and Paraseq 0.4.8 [21]. Needletail is the fastest single-threaded library to our knowledge, while Paraseq is getting increasingly popular because of its parallel-friendly design. The benchmarks presented in this section were run using the commit `5557b93` of our library. We measured the throughput of each parser in three main scenarios: parsing a genome in multiline FASTA format, short reads in FASTQ format and long reads in FASTQ format. The datasets used are detailed in Table 1.

**Table 1.**
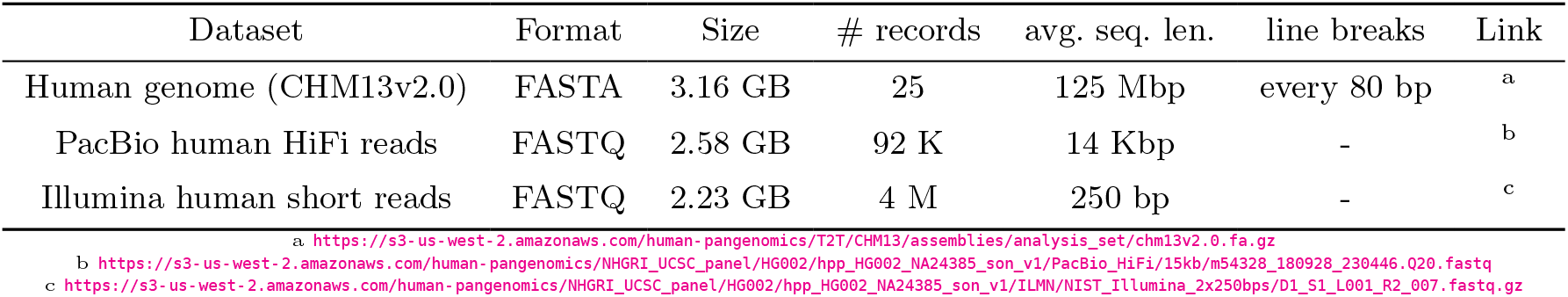
Description of the datasets benchmarked in the article.

The benchmarks are orchestrated through a bash script^4^, which iterates over all input datasets, runs the parsers in release mode, and records the results in a CSV file. The compilation is configured to target the native microarchitecture of each machine, ensuring that the generated code fully exploits the available CPU features. Multiple parser configurations are evaluated to capture performance sensitivity to instruction-level differences. Each measurement is averaged over ten repetitions.

To ensure representative and robust measurements, the benchmarks were deployed across a wide range of heterogeneous machines available on Grid’5000 [1] (with different CPU generations and microarchitectures). Jobs were executed on reserved nodes in isolation from other workloads, ensuring that no external interference perturbs the results. This setup provides a broad and controlled view of performance across architectures. For completeness, the same benchmarks were also run on the author’s personal laptop and desktop systems, allowing comparison with more conventional environments. A description of all the CPUs used across the benchmarks can be found in Table 2 in appendix.

### 3.1 Parsing throughput

The main metric that we evaluated is the throughput of each parser on the different datasets. In order to have a fair comparison that reflects a typical execution, the performance was measured when reading the files from disk and are thus subject to more significant I/O bottlenecks than for data loaded in RAM. Figure 1 shows the throughput when parsing a human genome (in FASTA format) and short reads (in FASTQ format). An additional plot for long reads is available in Figure 3 in appendix. For completeness, we also included results on data loaded in RAM in Figure 4 and results comparing the throughput of collecting the sequence string and only counting the number of bases in Figure 5 in appendix.

**Fig. 1.**
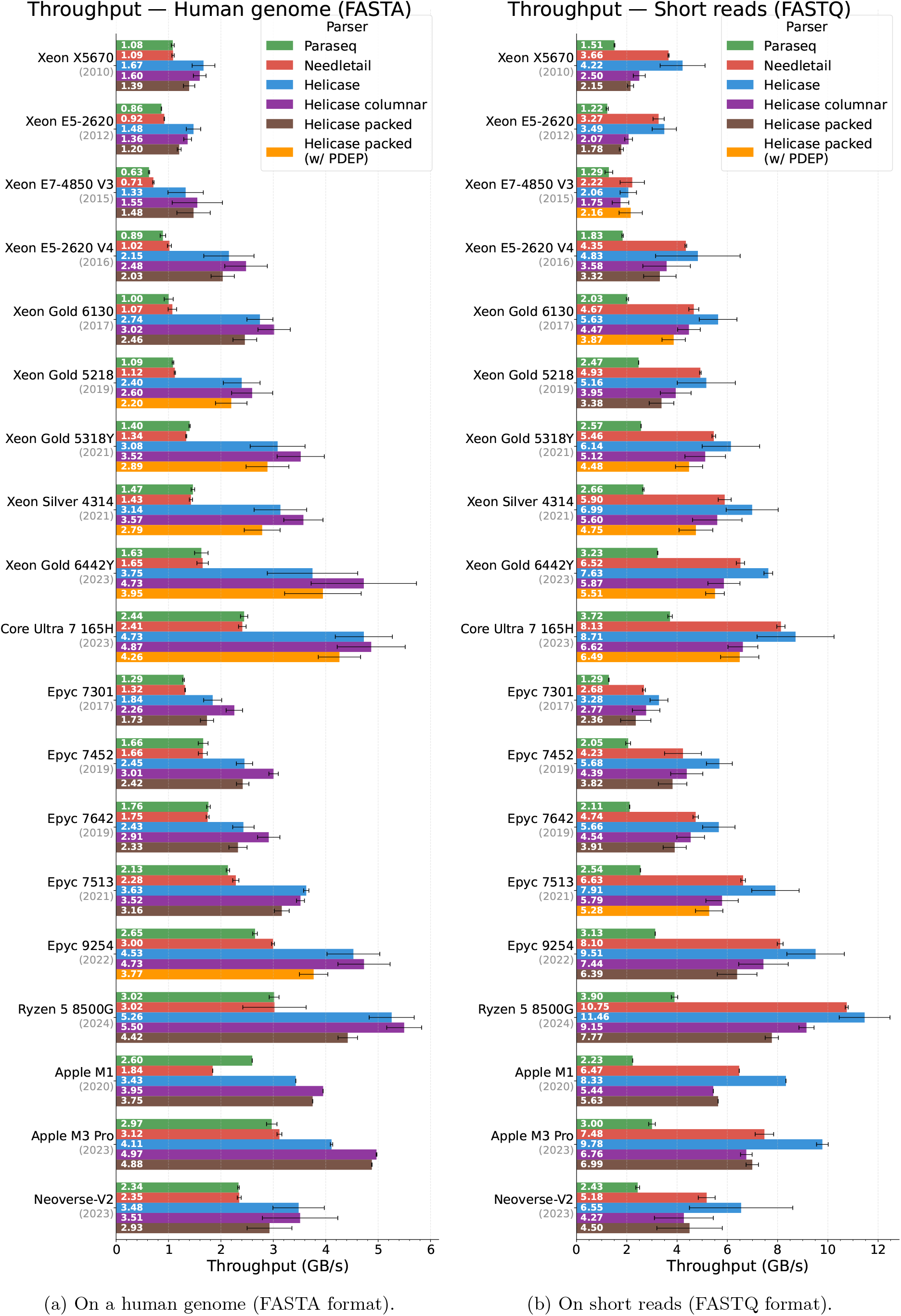
Throughput of each parser for FASTA and FASTQ files read from disk on multiple CPUs, sorted by manufacturer and year.

When parsing the human genome, Helicase is consistently twice as fast as its competitors on Intel CPUs, and roughly 50% faster on AMD and ARM CPUs. This trend is confirmed over all generations of CPUs, from the oldest ones only supporting SSE to those supporting AVX2 or NEON. We also observe that both bitpacking strategies (columnar and packed) are competitive with plain string parsing in this setting, which can be explained by the fact that the string version still has to eliminate the newlines contained in the sequence and thus cannot be copy-free. As expected given the extra interleaving step it requires, computing the packed representation is always slightly slower than the columnar one. Our implementation scales better with newer hardware. On old CPUs (e.g. Xeon X5670, 2010), the throughput is around 1.1 GB/s for Needletail and Paraseq and between 1.4 and 1.6 for variants of Helicase. Meanwhile, in the latest generation (Xeon Gold 6442Y, 2023), the throughput for Needletail and Paraseq ranges between 1.3 and 1.4 GB/s while Helicase achieves a throughput ranging from 3.7 to 4.7 GB/s. Hence, and contrary to the baseline, the implementation of Helicase clearly leverages hardware improvements of the last decade.

When parsing short reads, Helicase is consistently 10-30% faster than Needletail, itself faster than Paraseq. Compared to the results on the human genome, we observe that the overall throughput is higher and that the difference between Helicase and Needletail is less pronounced. This is mainly explained because FASTQ parsing requires less work (keeping track of the line count is sufficient to transition between states) and because the sequence of each record is not split across multiple lines, thus allowing a zero-copy strategy. On the other hand, bitpacked configurations do not benefit from this property and are comparatively slower than outputting plain strings. The results on long reads, shown in Figure 3 in appendix, are even less pronounced. Having longer contiguous sequences make the zero-copy strategy is even more noticeable and file I/O becomes a bigger limitation.

To illustrate the impact of compile-time specialization of the parser, we compared the throughput of collecting the sequence string for each record against a configuration that only counts the number of ACTG bases (skipping other IUPAC codes) on the human genome. This comparison, presented in Figure 5 in appendix, confirms that a lighter configuration requirement successfully produces more efficient code.

### 3.2 Number of instructions, cycles and branches per byte

In addition to throughput, we measured the number of instructions, cycles and branches corresponding to each execution using perf counters, and divided it by the input size to have uniform values. Since perf counters are not as widely available on macOS, we did not include Apple CPUs for these metrics. Figure 2 shows the number of instructions and cycles per bytes for each parser on the human genome. Additional plots on short reads are available in Figure 6 in appendix, as well as plots on the number of branches per byte in Figure 7 and the number of branch misses per MB in Figure 8.

**Fig. 2.**
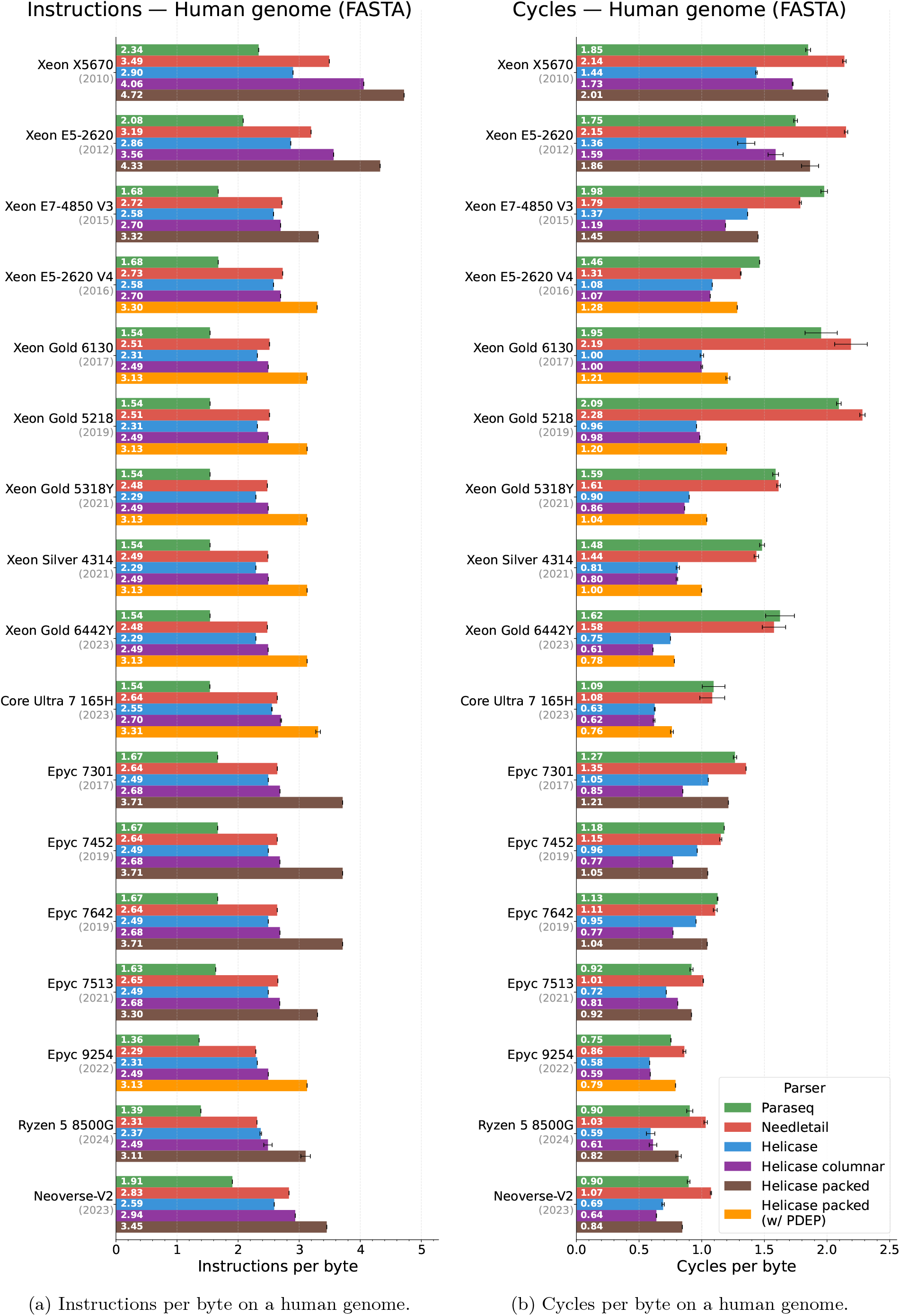
Instructions and cycles per byte on multiple CPUs, sorted by manufacturer and year.

These plots provide many interesting insights on the CPU characteristics and how well each parser exploits them. First of all, we note that the main factor impacting the number of instructions is the instruction-set supported by each CPU. This is especially noticeable when comparing Intel CPUs only supporting SSE (first two Xeon CPUs in Figure 2a, prior to 2013) to those supporting AVX2 afterwards: since SSE vectors are twice as small as AVX vectors, scanning a given block requires more instructions. When focusing on the packed representation, we can also observe the shift from microcoded PDEP instructions (first three Epyc CPUs, prior to 2020), which we avoided in favor of a fallback implementation shown in brown, to native PDEP instructions on AMD CPUs over the years.

Even though the number of instructions is roughly constant for a given instruction-set, the number of cycles spent to execute them is not. We observe in Figure 2b a decrease in the number of cycles over the years, reflecting the continuous improvements of CPU manufacturers. However, we can also notice that while Helicase has a number of instructions similar to Needletail, it uses significantly less cycles. This suggests that our implementation relies on instructions that the CPUs evaluate in parallel, making a better use of vectorized instructions, and therefore allows a good pipelining strategy. Finally, the results on the number of branches (shown in Figure 7 in appendix) confirm that Helicase successfully eliminates more branches with our vectorization strategy.

## 4 Conclusion

We presented Helicase, a high-throughput Rust library for parsing FASTA and FASTQ files that exploits SIMD vectorization to maximize single-threaded throughput on both x86 and ARM. At the core of our approach is a vectorized lexing stage based on bitmask classifiers derived from the theory of counter-free automata [13,19]. Rather than scanning the input byte by byte, the lexer annotates 64-byte blocks in parallel using carry-propagating arithmetic and bitwise operations, fusing the detection of all structural markers into a single pass. The parser is then structured as a finite state machine over these bitmasks, and specialized at compile time to only contain the code paths relevant to the requested fields. Helicase is able to produce on-the-fly DNA bitpacking for two compact representations: a *packed* format and a *columnar* format that separates the high and low bits, with non-ACTG characters handled either as segment boundaries or via a lossy encoding with an additional ambiguity mask. We also interfaced Helicase with the existing libraries developed for SimdMinimizers [5], to support efficient *k*-mer hashing and minimizer computation for the packed format.

We evaluated Helicase on a wide collection of CPUs, deployed reproducibly on Grid’5000. On FASTA genome parsing, Helicase is up to 2 × faster than Needletail on Intel CPUs and 50% faster on AMD and ARM, while matching or exceeding it on all FASTQ workloads. For data loaded in RAM, Helicase reaches the core memory bandwidth (49 GB/s on an Apple M3 Pro for long reads), confirming that the vectorized design successfully eliminates the parsing overhead as a bottleneck.

A number of avenues remain to extend and improve our approach. On the memory management side, DNA records are currently stored entirely in main memory, which is sufficient for most organisms but may fall short for certain species whose chromosome sizes exceed human genome standards by an order of magnitude [18]. As such large genomes accumulate through ambitious sequencing programs [16], introducing a configurable buffer size to process DNA records under memory constraints is a natural next step. On the hardware side, implementing a version optimized for AVX-512 could provide substantial performance gains on latest CPUs. Parallelization is another important direction: since Helicase processes a file as a single stream, one could instead partition the input into multiple chunks and assign each to a separate reader, avoiding the synchronization overhead of a producer-consumer model. On the algorithmic side, there is potential to further exploit the columnar DNA representation by designing specialized algorithms for *k*-mer generation and hashing or pattern matching.

In addition to FASTA and FASTQ, Helicase could also be extended to parse annotations in GFF and GTF formats efficiently. Another extension involves packaging Helicase for interpreted languages such as Python and JavaScript. While this would broaden accessibility, it introduces challenges related to managing the configuration state and controlling runtime code size. Finally, building additional tooling on top of the core library, such as utilities for format conversion, filtering or statistics, would further integrate Helicase into everyday workflows.

## Acknowledgments

This work is funded by the French National Research Agency SxC ANR-24-CE25-2874-01, and has benefited from funding from the French State under the France 2030 program, reference ANR-21-IDES-0006. The European Metropolis of Lille and the University of Lille are also acknowledged for their funding and support of the project WILL-CHAIRES-25-001-BOSSA. The authors would like to thank Lucas Robidou for proofreading, Ragnar Groot Koerkamp for insightful discussions and Marius Bilasco for running some of the benchmarks.

## A Features of the benchmarked CPUs

**Table 2.** Instruction-set extensions supported by the benchmarked CPUs.

| CPU | Year | SSE | AVX2 | BMI2 | NEON |
| --- | --- | --- | --- | --- | --- |
| Intel Xeon X5670 | 2010 | ✓ | – | – | – |
| Intel Xeon E5-2620 | 2012 | ✓ | – | – | – |
| Intel Xeon E7-4850 V3 | 2015 | ✓ | ✓ | ✓ | – |
| Intel Xeon E5-2620 V4 | 2016 | ✓ | ✓ | ✓ | – |
| Intel Xeon Gold 6130 | 2017 | ✓ | ✓ | ✓ | – |
| Intel Xeon Gold 5218 | 2019 | ✓ | ✓ | ✓ | – |
| Intel Xeon Gold 5318Y | 2021 | ✓ | ✓ | ✓ | – |
| Intel Xeon Silver 4314 | 2021 | ✓ | ✓ | ✓ | – |
| Intel Xeon Gold 6442Y | 2023 | ✓ | ✓ | ✓ | – |
| Intel Core Ultra 7 165H * | 2023 | ✓ | ✓ | ✓ | – |
| AMD Epyc 7301 | 2017 | ✓ | ✓ | ~ | – |
| AMD Epyc 7452 | 2019 | ✓ | ✓ | ~ | – |
| AMD Epyc 7642 | 2019 | ✓ | ✓ | ~ | – |
| AMD Epyc 7513 | 2021 | ✓ | ✓ | ✓ | – |
| AMD Epyc 9254 | 2022 | ✓ | ✓ | ✓ | – |
| AMD Ryzen 5 8500G * | 2024 | ✓ | ✓ | ✓ | – |
| Apple M1 * | 2020 | – | – | – | ✓ |
| Apple M3 * | 2023 | – | – | – | ✓ |
| ARM Neoverse-V2 | 2023 | – | – | – | ✓ |

~ microcoded PDEP/PEXT instructions (very slow)
\*personal computers, not benchmarked in a reproducible environment

## B Additional figures

### B.1 Additional experiments on data read from disk

**Fig. 3.**
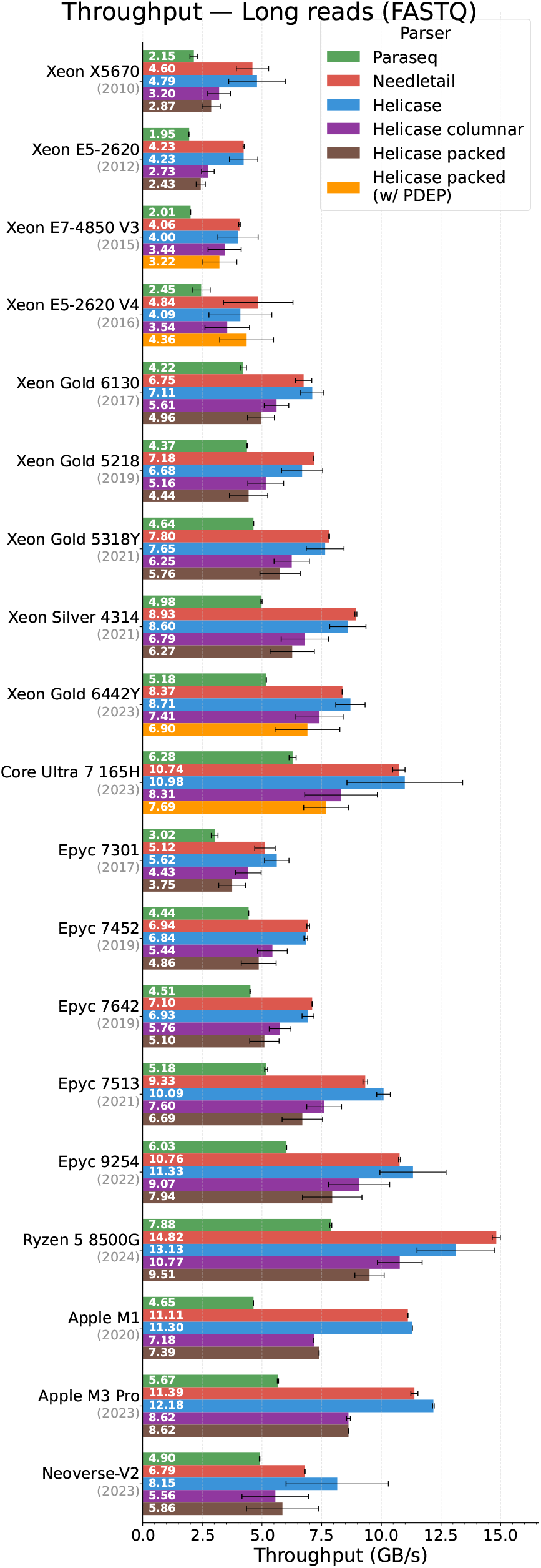
Throughput of each parser for long reads on multiple CPUs, sorted by manufacturer and year.

### B.2 Additional experiments on data loaded in RAM

**Fig. 4.**
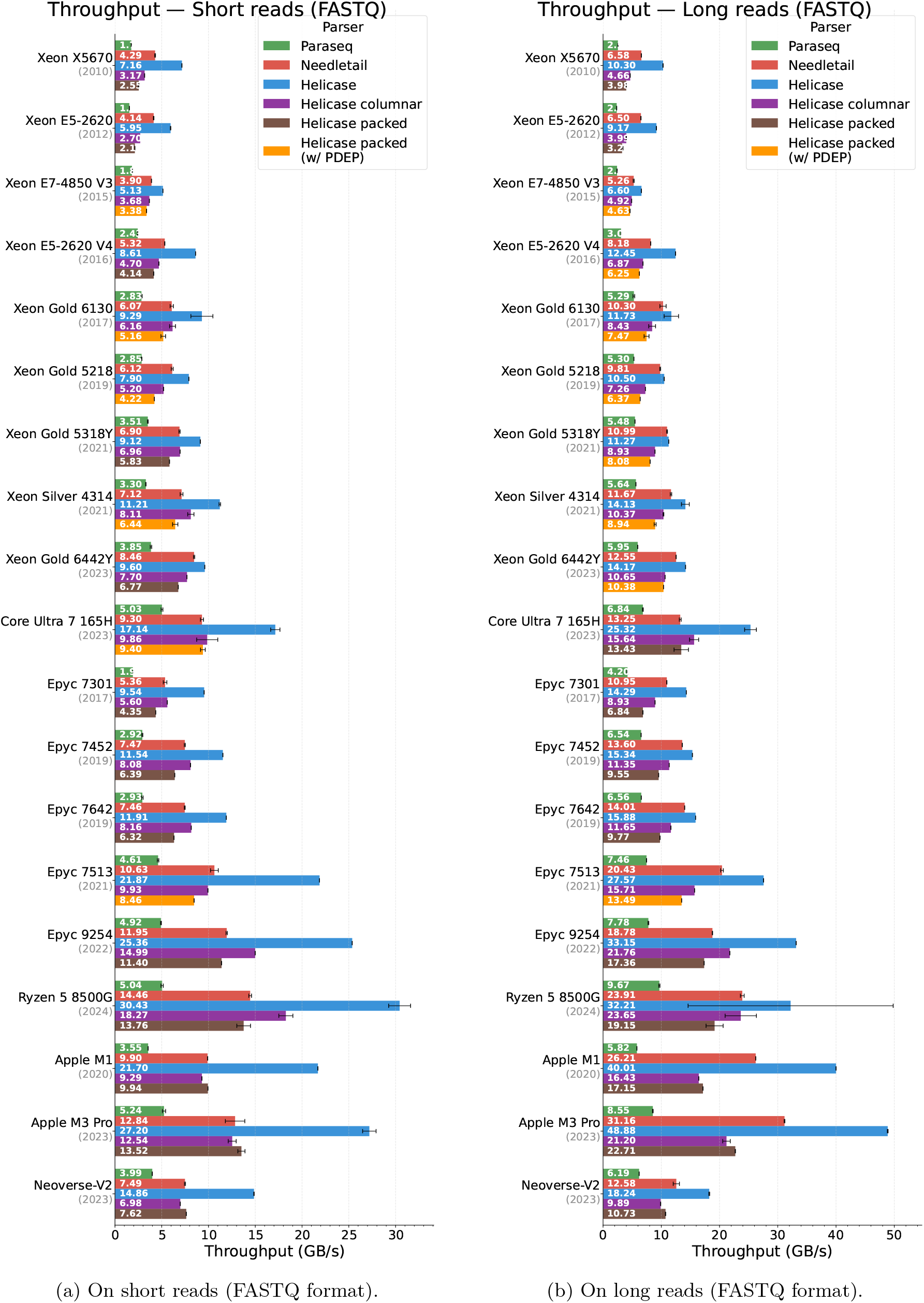
Throughput of each parser for data loaded in RAM on multiple CPUs, sorted by manufacturer and year. Needletail and Paraseq both have to use a reader over a slice, which degrades their performance.

**Fig. 5.**
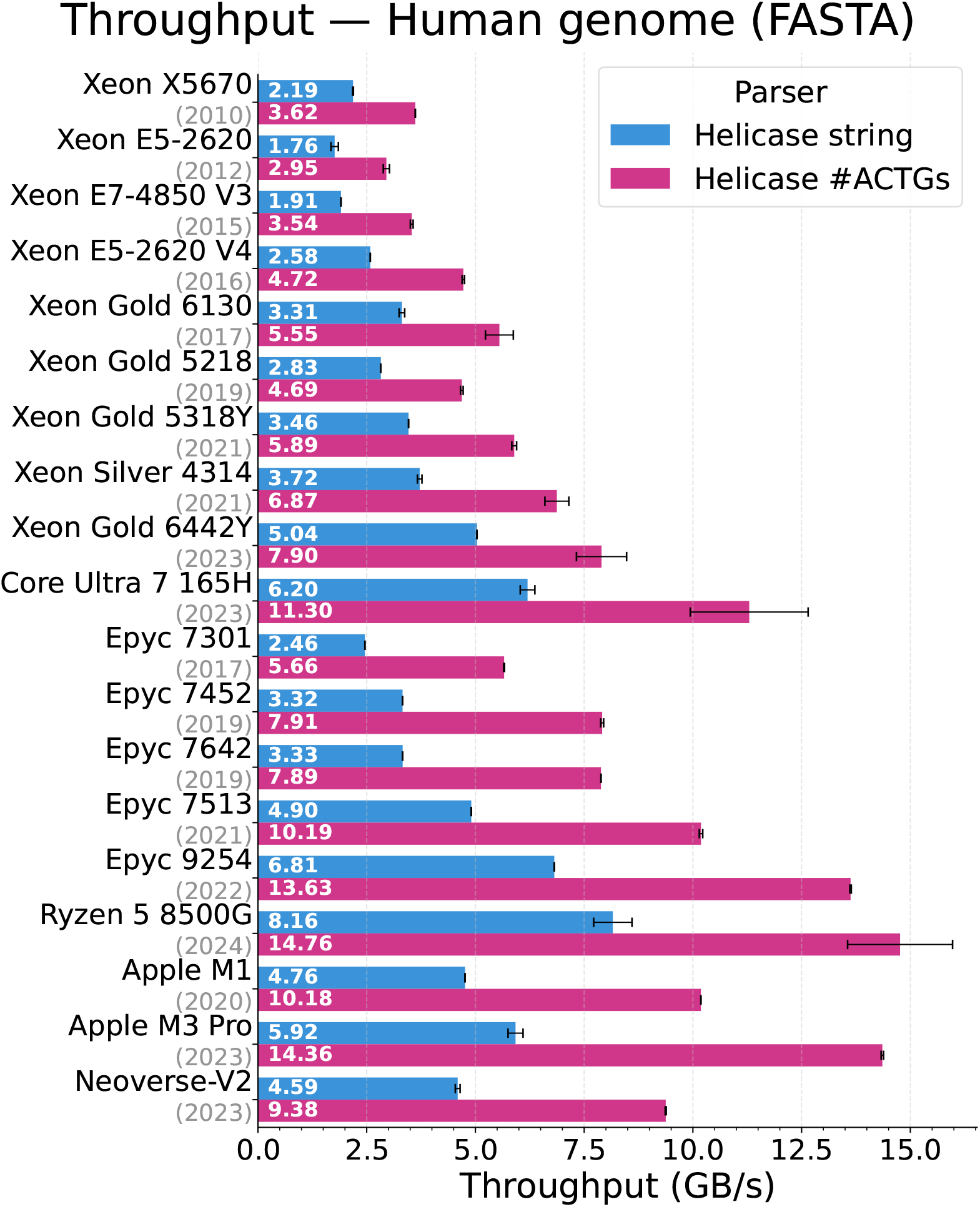
Throughput of Helicase string collection compared to counting DNA bases.

**Fig. 6.**
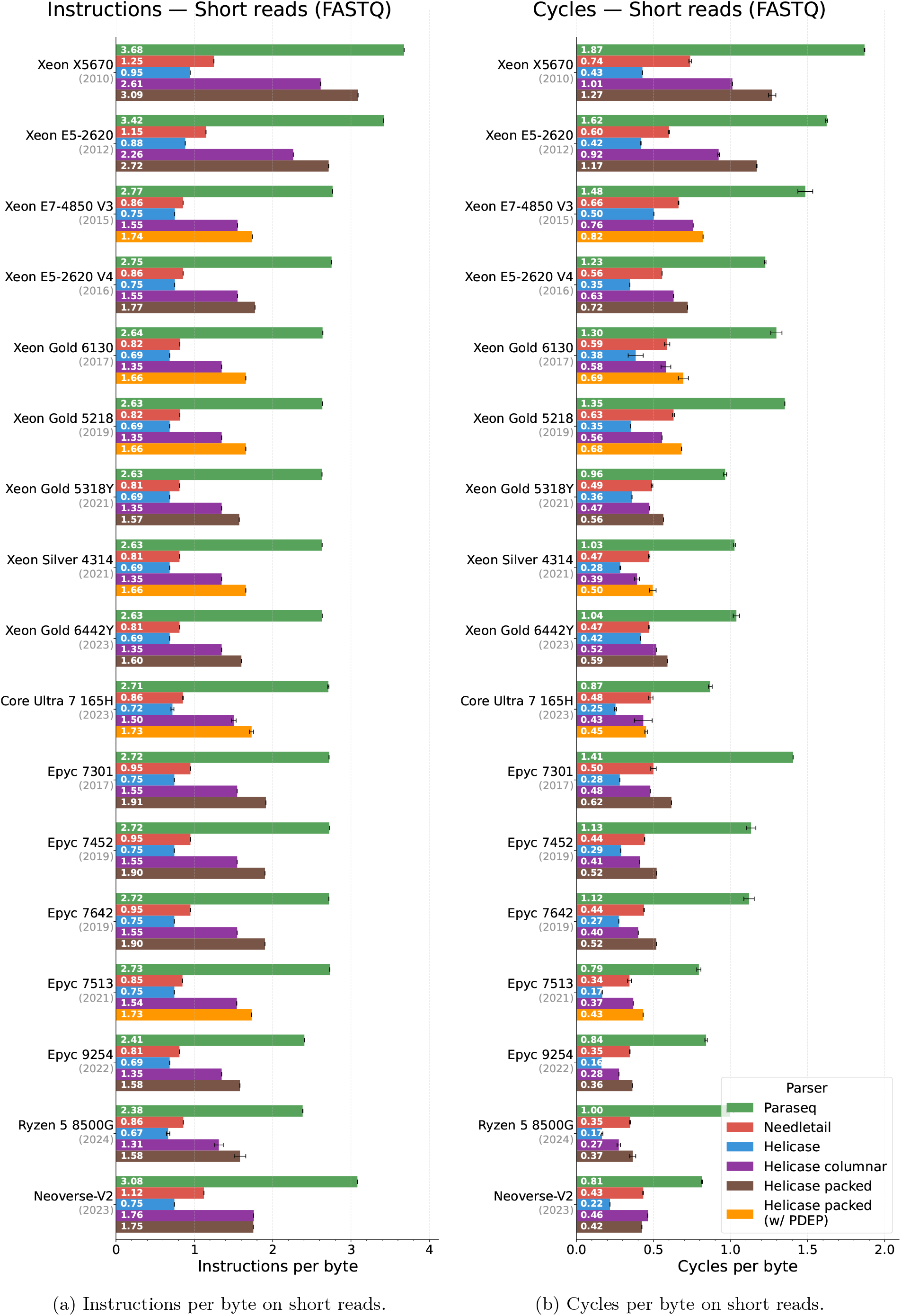
Instructions and cycles per byte on multiple CPUs, sorted by manufacturer and year.

**Fig. 7.**
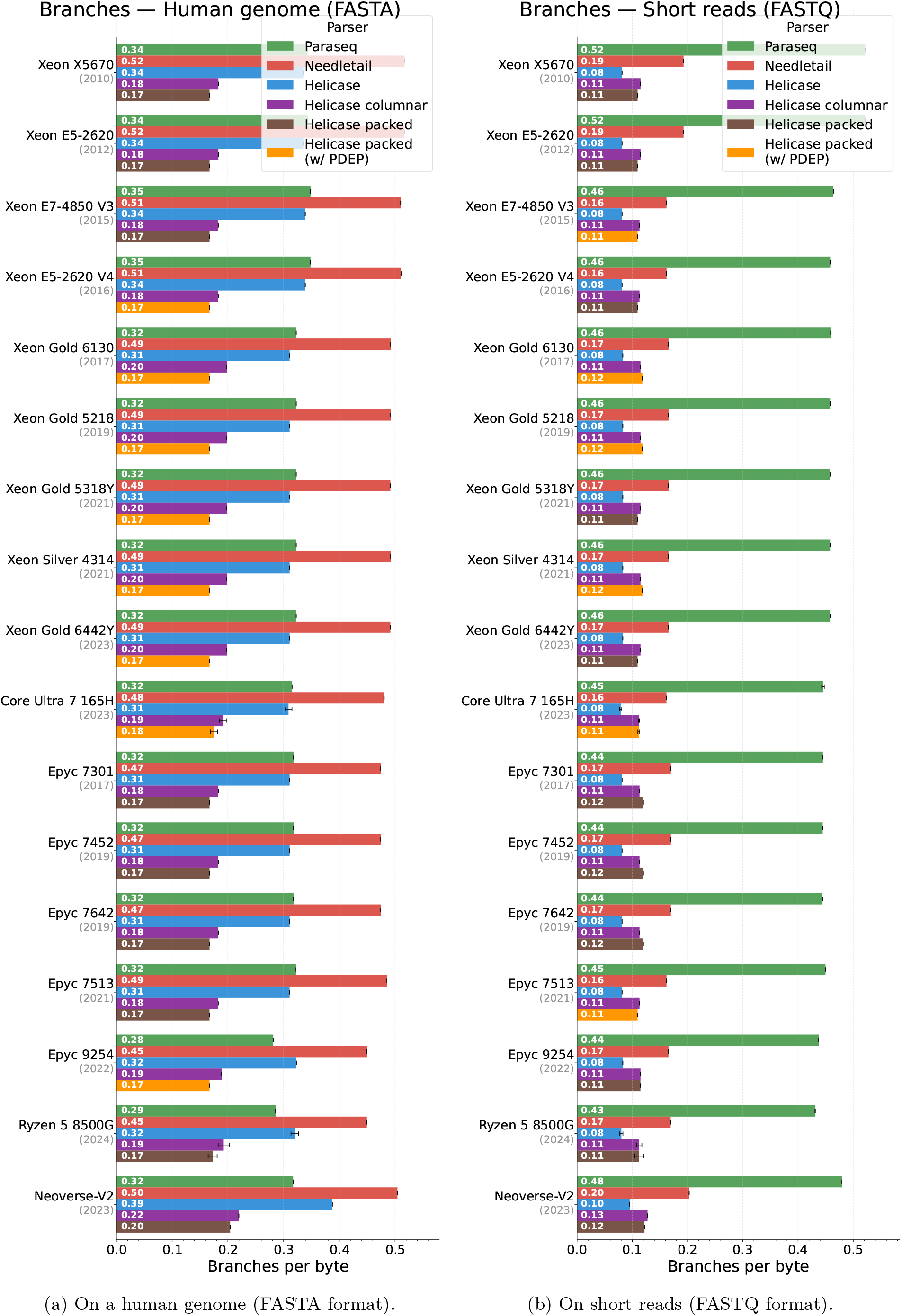
Branches per byte on multiple CPUs, sorted by manufacturer and year.

**Fig. 8.**
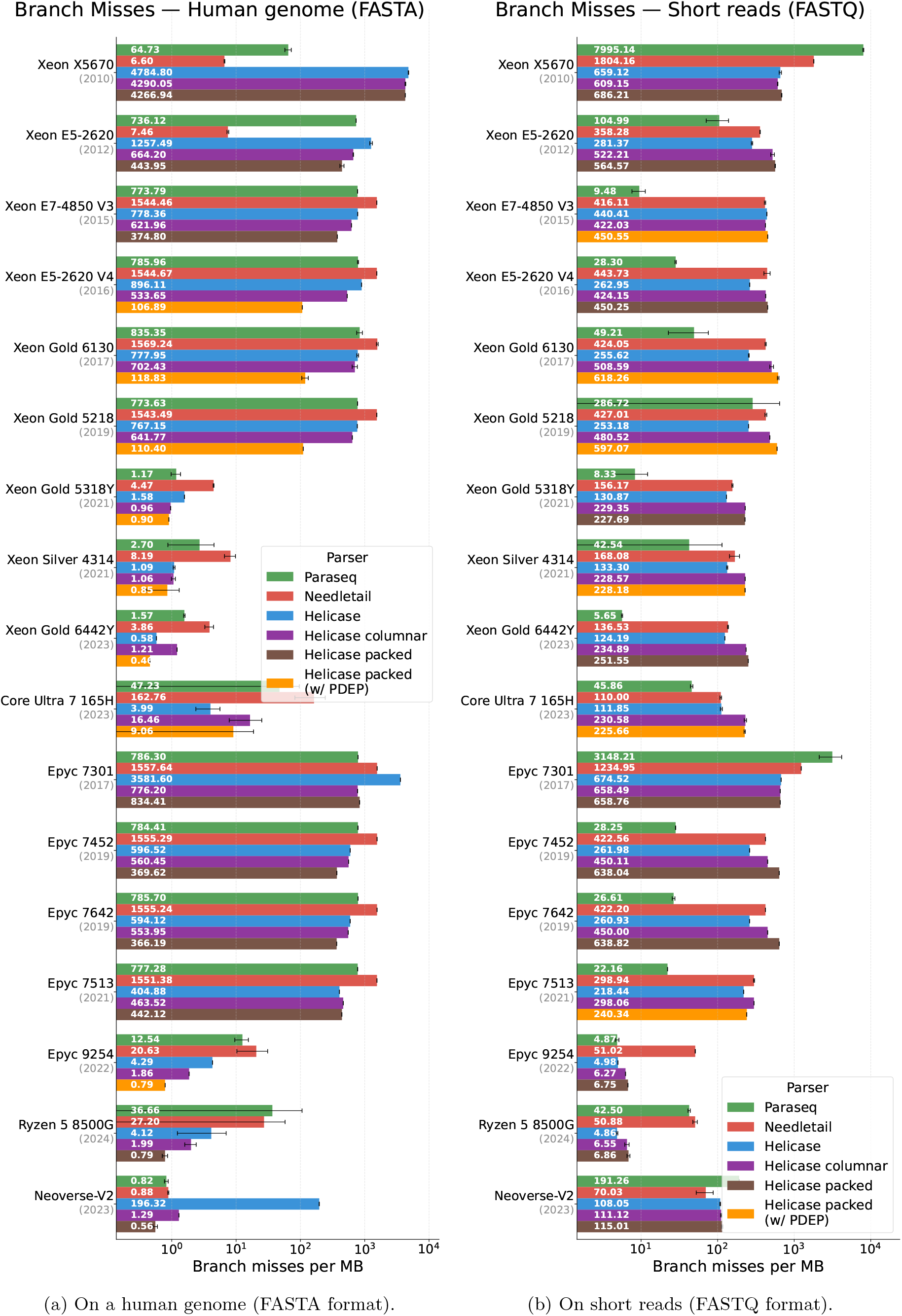
Branch misses per MB on multiple CPUs, sorted by manufacturer and year.

## Footnotes

3 https://stackoverflow.com/questions/74722950

4 https://github.com/imartayan/helicase/blob/main/bench/bench.sh

